# Anodal Transcranial Direct Current Stimulation (tDCS) of hMT+ Does Not Affect Motion Perception Learning

**DOI:** 10.1101/405696

**Authors:** Stephanie J. Larcombe, Christopher Kennard, Jacinta O’Shea, Holly Bridge

## Abstract

**Background:** Human visual cortical area hMT+, like its homologue MT in the macaque monkey, has been shown to be particularly selective to visual motion. After damage to the primary visual cortex (V1), patients often exhibit preserved ability to detect moving stimuli, which is associated with neural activity in area hMT+. As an anatomical substrate underlying residual function in the absence of V1, promoting functional plasticity in hMT+ could potentially boost visual performance despite cortical damage.

**Objective:** To establish in healthy participants whether it is possible to use transcranial direct current stimulation (tDCS) over hMT+ to potentiate learning of visual motion direction discrimination.

**Methods:** Participants were trained daily for five days on a visual motion direction discrimination task. Task difficulty was increased as performance improved, by decreasing the proportion of coherently moving dots, such that participants were always performing at psychophysical threshold. tDCS, either anodal or sham, was applied daily during the 20-minute training session. Task performance was assessed at baseline and at the end of the training period.

**Results:** All participants showed improved task performance both during and after training. Contrary to our hypothesis, anodal tDCS did not further improve performance compared to sham stimulation. Bayesian statistics indicated significant evidence in favour of the null hypothesis.

**Conclusion:** Anodal tDCS to hMT+ does not enhance visual motion direction discrimination learning in the young healthy visual system.

## Introduction

The principal pathway conveying visual information from the eye to the brain projects via the primary visual cortex (V1), the largest cortical visual area. The critical role of this area in vision is reflected in the fact that any damage to this region can lead to cortical blindness. However, even after damage to V1, many patients continue to show cortical brain activity in the human motion area hMT+ [1-4] and some are adept at detecting moving stimuli, a capacity known as blindsight [5]. Hence area hMT+ is a potential intervention target for rehabilitation regimes that aim to improve visual function after V1 damage [6, 7].

In the healthy visual system, the specialised role of hMT+ in humans and MT in the non-human primate has been demonstrated using multiple techniques, including electrophysiology[8-10], lesion studies [11-13], fMRI [14] and electrical stimulation [15]. Given this role it could be hypothesized that perceptual training on motion discrimination should result in functional changes within MT. However, this does not appear to be the case, at least in the macaque. Law & Gold [16-18] have shown that learning a motion task does not change neuronal properties in MT, but rather this occurs at the level of the sensory-motor decision, in lateral intraparietal area (LIP). Nevertheless, Lui & Pack [19] demonstrated that while training on a motion discrimination task did not change the sensitivity of individual MT neurons, after training there was an increased effect of MT microstimulation on biasing motion direction decisions.

In humans, learning a visual motion discrimination task over 5 days causes an increase in neural activity in MST, part of the human motion complex, which correlates with the amount of learning [20], suggesting a functional role for MST in the improved performance. Since this region often remains active in patients who have suffered damage to V1, it may be that visual discrimination training could strengthen subcortical connections to visual motion areas and increase residual visual function. While boosting performance with training is beneficial, addition of an adjunct intervention to increase plasticity, such as pharmacological enhancement of acetylcholine levels [21], can further potentiate the effect.

Here we tested whether a different neuroplasticity intervention, non-invasive brain stimulation of hMT+, when applied during training could also increase learning. We chose to stimulate using excitatory (anodal) transcranial direct current stimulation (tDCS) and compare this to sham. Anodal tDCS increases visual cortical excitability [22, 23] and has been reported to enhance visual functioning [24-27]. In the motor system, anodal tDCS applied to primary motor cortex during training has been shown to enhance acquisition and consolidation of motor learning [28, 29]. The current study tested whether anodal tDCS of hMT+ would augment learning of visual motion direction discrimination.

## Materials and Methods

### Participants

24 participants (13 female and 11 male; M=24.7 years; SD=5.8 years) were randomly assigned to an anodal (n=13) or sham (n=11) stimulation group. Before study completion, three participants withdrew from the study, two from the anodal group and one from the sham group. Owing to incomplete data, these participants were excluded from all analyses.

The study was approved by the local InterDivisional Research Ethics Committee (IDREC) at the University of Oxford (reference MSD-IDREC-C2-2014-025) and all participants gave written, informed consent. Research was carried out in accordance with the Code of Ethics of the World Medical Association (Declaration of Helsinki). All participants underwent safety screening to exclude contraindications to brain stimulation prior to each test session.

### Visual task

Participants completed a motion perception task where the instructions were to discriminate the direction of coherently-moving dots presented amongst randomly-moving distractor dots. Moving dots (n=143) were presented within a circular area 11° in diameter, offset 10° to the left or right of fixation. Dots were high contrast white dots on a black background. The luminance and chromaticity measures (SpectraScan PR-650) were white: 96.8cd/m^2^ (x=0.289, y=0.312), and black: 0.92cd/m^2^ (x=0.236, y=0.247). Each trial consisted of a 500ms stimulus window, a pause for the participant response, and a 200ms feedback window (Figure 1A). The next trial began automatically following the feedback window. The response window remained on-screen until the participant responded. During all sessions participants were offered an optional screen break every 20 trials to reduce fatigue.

**Figure 1:**
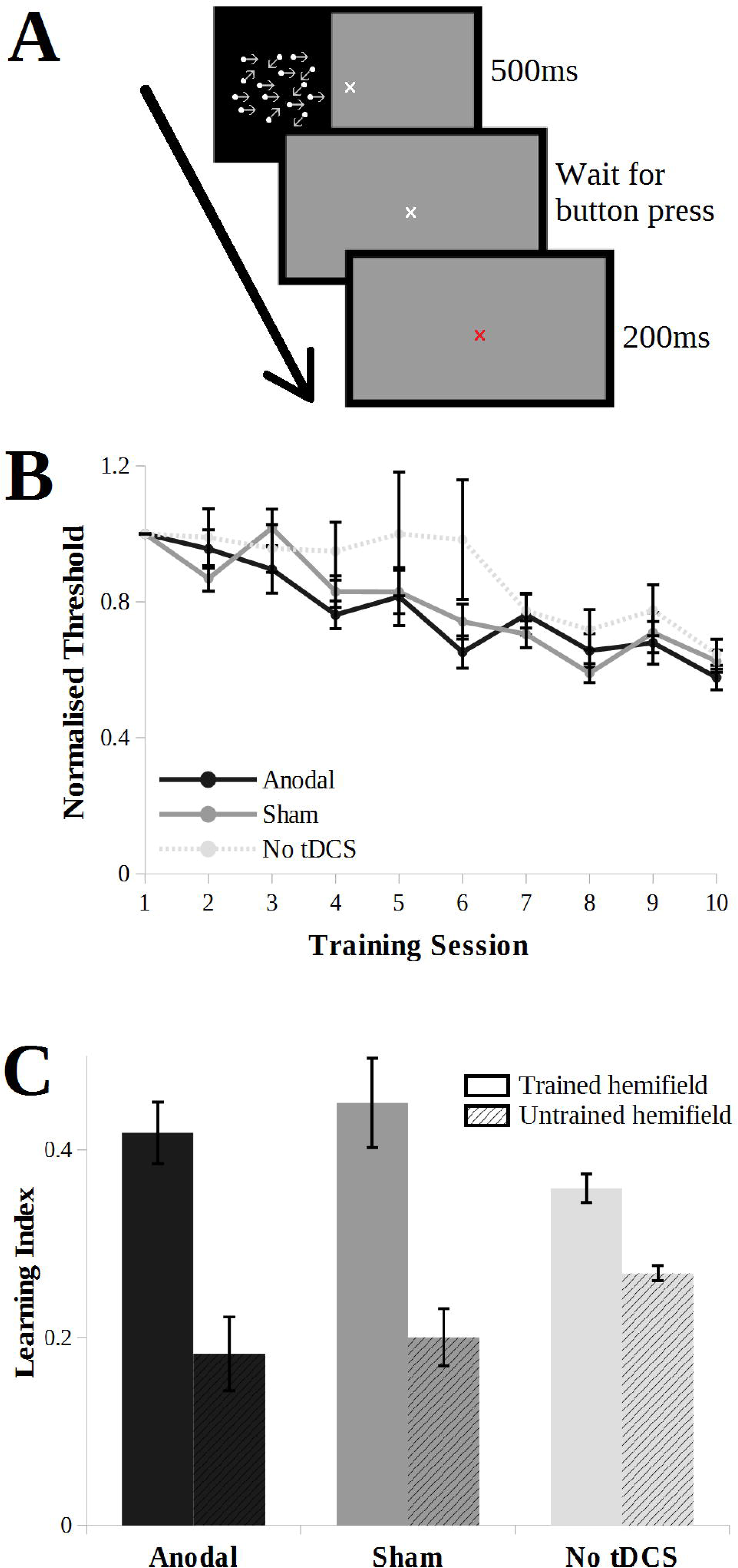
A: Motion direction discrimination task. Participants determined the direction of coherent motion of moving dots. Each trial consisted of 500ms stimulus period, followed by an untimed user response window. Following participant response, feedback was provided (red or green fixation cross) for 200ms, and then the next trial start immediately. **B: Comparison of performance of anodal, sham and no tDCS stimulation groups across the ten training sessions.** There was no significant difference between normalised discrimination thresholds at the final training session, indicating a similar amount of learning occurred in all three groups. **C: Comparison of learning index computed from the assessment on day 1 and day 5 of anodal, sham and no tDCS stimulation groups across trained and untrained visual hemifields.** There was no significant difference in the learning index across groups, neither in the trained nor untrained hemifields. Error bars show ±SEM.

All participants completed ten training sessions of the motion discrimination task. The training sessions were completed two per day, for five consecutive days (2 training sessions of 400 trials per day, each session lasting around 10 minutes) with a break of 1-2 minutes between training sessions carried out on the same day. Learning effect was quantified from the assessment sessions on day 1 and day 5, which acted as the dependent variable (400 trials per assessment, each session lasting around 20 minutes). In these assessment sessions, stimuli were presented to the left or right visual hemifield in a pseudorandomly interleaved manner, with 200 trials per hemifield. For the training sessions, the stimulus was delivered to the right visual hemifield only, to allow the left hemifield to act as a control (i.e. contrast trained>untrained hemifield).

Task difficulty was adaptively modulated by altering the ratio of coherently-moving dots to randomly-moving dots, using a two up one down staircase procedure[30]. New staircases were initiated for every assessment and training session. For the assessment sessions, independent staircases were applied for the two visual hemifields. Motion direction discrimination thresholds for every session were calculated by taking the mean of the coherence on each reversal trial (the task changed from increasing in difficulty to decreasing, or vice versa). The first 10 reversals were discarded. The average provided a threshold at which the participant is predicted statistically to be correct 80% of the time.

### Brain stimulation

Participants received five sessions (20 minutes each) of tDCS delivered over left hMT+ (HDCkit, Magstim), one each day, concurrent with the 20-minute training period. For sham stimulation the current was ramped up to 1mA over 10 seconds and then switched off. For anodal stimulation, the current was ramped up over a duration of 10 seconds and remained at 1mA for 20 minutes. Direct current was delivered through electrodes inside rectangular saline-soaked sponges. The cathode (8.5cm x 6cm) was placed at the vertex and the anode (5cm x 5cm) was placed 3cm above the inion along the nasion-inion line and 6cm left of the midline in the sagittal plane (Figure 1). The latter scalp coordinates were derived from prior research with transcranial magnetic stimulation (TMS), which showed effects of stimulation at this location on visual motion processing [31, 32]. The electrode montage used here has been used in previous tDCS research to stimulate left hMT+ [26].

The experimenter who conducted the training and stimulation was blinded as to whether the participant was receiving sham or anodal stimulation. This was done using an automatic blinding mode on the tDCS stimulation device. Unblinding was performed once data collection was completed, prior to analysis.

## Results

There were no reported adverse effects of the tDCS, with the exception of sensations of itching and tingling in both sham and anodal groups, with no difference between the groups.

Data from ten participants in a previous study (5 female, 18-29 years) using the same protocol, but without any stimulation, were included in the analysis for comparison [33]. For all assessment and training sessions, performance was quantified by determining the motion direction discrimination threshold, a measure used in previous studies to quantify changes in learning [20, 33].

For training sessions, thresholds were normalized within each participant relative to performance in the initial training session (i.e. Day 1). Performance levels in the daily motion perception training sessions were indistinguishable across the three groups (Figure 1B). While there was a significant effect of training session (F(9,252) = 16.3; p < 0.001), indicating that participants learned the task, there was no difference between the anodal, sham and no stimulation groups (F(2, 28) = 1.5; p = 0.23).

To aid interpretation of the null effect of tDCS, a Bayesian repeated measures ANOVA was also performed, using the open-source software package JASP (http://www.jasp-stats.org) [34]. Bayesian analyses permit a test of the relative strength of evidence for the null hypothesis (H_0_: no effect of tDCS stimulation group) versus the alternative hypothesis (H_1_: change in behaviour as a result of tDCS condition) [35]. The pattern of results was consistent across both frequentist and Bayesian analyses. The main effect of training was significant, reflected in a higher Bayes factor for the alternative hypothesis (H_1_: training changes behavioural performance) than the null hypothesis (H_0_: no behavioural effect of training; BF_10_ = 1.6 × 10^18^). The Bayes factor for the effect of tDCS stimulation condition (H_1_) was less than one (BF_10_ = 0.37). In contrast, the reciprocal value (BF_01_ = 2.7) suggests that the null hypothesis (that there is no effect of tDCS condition) is 2.7 times more likely than the alternative hypothesis.

For assessment sessions, a learning index was calculated using the following formula:

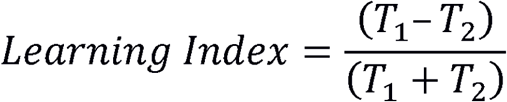

where T_1_ refers to the threshold before training, and T_2_ refers to the threshold after training was completed.

There was no significant difference in the learning index between anodal, sham and no stimulation groups (Figure 1C) either for the trained hemifield (one-way ANOVA: F(2, 30)=1.754, p=0.192) or the untrained hemifield (one-way ANOVA: F(2, 30)=2.283, p=0.121). A one-way Bayesian ANOVA was also performed on the learning index, and, consistent with the previous result, provided evidence in favour of the null hypothesis (BF_10_ = 0.63; BF_01_ = 1.59).

Next we tested if anodal tDCS would enhance consolidation of visual learning across consecutive days. Offline consolidation refers to performance gains that occur after training during a rest interval. In this task, offline consolidation would be reflected in a lower direction discrimination threshold the day after training compared to the threshold achieved at the end of the previous day. Forgetting would be reflected in a threshold increase. Maintenance of learning would be reflected in no change across the interval between days. Figure 1B indicates there was no clear evidence of offline consolidation across consecutive days. A one-way ANOVA on the mean difference in performance between consecutive days indicated no effect of tDCS on consolication (F(2,30 = 1.52, p = 0.24). Similarly, the Bayesian ANOVA provided evidence in favour of the null hypothesis (BF_10_ = 0.54; BF_01_ = 1.85).

## Discussion

All participant groups included in this study showed significant improvement in direction discrimination thresholds over the five-day training period, consistent with previous results [20, 33]. Furthermore, daily anodal tDCS to hMT+ during training had no effect on learning or offline consolidation.

All groups showed improved thresholds, i.e. learned from training. Yet, despite using stimulation parameters closely similar to previous tDCS studies of hMT+ [36], there was no difference in performance between groups receiving anodal or sham tDCS. The improvement with training in both these groups was comparable to previous data from participants that had not received stimulation (Figure 1C). There are several potential reasons for the lack of a tDCS learning or consolidation enhancement effect.

Firstly, hMT+ was not identified in each participant individually using fMRI, so it is possible that the anodal electrode did not effectively stimulate the target area. However, this seems unlikely. Area hMT+ has been shown to vary only by approximately 2.7cm in the left hemisphere [37] and to be, on average, 0.3cm^3^ in size [38]. The tDCS electrode dimensions exceed this (hMT+ anode: 5cm × 5cm), so it is likely that the stimulation at least partially covered hMT+. A related point is that the stimulation is applied at the scalp, and the achieved current dose within cortex is likely to vary across participants. The location of hMT+ is also variable across individuals, and can be either on a gyrus, in the sulcus, or both [39, 40]. How variations in individual anatomy interact with induced electrical current dose is currently under active investigation [41]. Nevertheless, inter-participant variance was in a similar range for the anodal and sham groups, suggesting this is unlikely to be a key factor in the null result.

Secondly, only the effects of anodal stimulation on a motion direction perception task were considered in this study. It may be interesting to investigate whether cathodal stimulation of hMT+ alters motion perception in this type of extended training protocol. Battaglini et al. [42] found that both anodal and cathodal stimulation improved performance on a visual motion discrimination task, although the authors suggest the improvement was due to different mechanisms. We chose to stimulate with anodal tDCS as this polarity of stimulation has most reliably been associated with learning gains, at least in the motor system.

A third point relates to the number of participants in the study. Variability in tDCS effects have led to calls for greatly increased sample sizes [43]. One important, relatively neglected point in this discussion is that the end goal of much neuromodulation research is therapeutic. Here our motive for investigating tDCS was to advance the long-term goal of improving visual function in individual patients. For this to be practical, tDCS effects need to be measurable reliably in small samples, such as the single-case and small group designs that reflect the real-world challenges of clinical neuropsychology research and practice [44]. A small, but statistically significant effect that requires large populations to detect is unlikely to have measurable benefit at an individual level.

Finally, multiple studies have shown that visual perceptual learning improves visual performance. We found no evidence that concurrent anodal tDCS to hMT+ accelerated perceptual learning or enhanced consolidation over a 5-day training period. It is possible that the training itself induced a ceiling effect in these young participants with a healthy visual system.

Although tDCS in these healthy participants did not improve visual motion discrimination, this does not rule out the possibility of a beneficial effect of the same intervention in a patient group. In healthy, sighted participants the main thalamocortical projection from the retina to V1 is intact. In contrast, patients with damage to the primary visual cortex must rely on other connections to convey retinal information to the visual cortex. Since these alternative connections are unlikely to be as strong as the V1 pathway, it may be that training this pathway concurrent with electrical stimulation in patients would have a measurable effect.

## Conclusion

In conclusion, anodal stimulation of hMT+ in healthy participants during motion perception training did not improve performance compared to sham stimulation. This suggests that online, anodal stimulation of hMT+ (at least with the montage, current strength, duration, and participant sample tested here) may not be an effective way to modulate motion perception learning.

## Author Contributions

Conception and design of the work (SL, CK, HB, JO’S), data collection (SL, JO’S), data analysis and interpretation (SL, HB), drafting the article (SL, HB, JO’S), critical revision of the article (SL, HB, JO’S), final approval of the version to be published (SL, CK, HB, JO’S).

## References

[1] Ajina S, Kennard C, Rees G, Bridge H. Motion area V5/MT+ response to global motion in the absence of V1 resembles early visual cortex. Brain 2015;138(Pt 1):164–78.

[2] Bridge H, Hicks SL, Xie J, Okell TW, Mannan S, Alexander I, et al. Visual activation of extra-striate cortex in the absence of V1 activation. Neuropsychologia 2010;48(14):4148–54.

[3] Morland AB, Le S, Carroll E, Hoffmann MB, Pambakian A. The role of spared calcarine cortex and lateral occipital cortex in the responses of human hemianopes to visual motion. J Cogn Neurosci 2004;16(2):204–18.

[4] Zeki S, Ffytche DH. The Riddoch syndrome: insights into the neurobiology of conscious vision. Brain 1998;121 (Pt 1):25–45.

[5] Cowey A. The blindsight saga. Exp Brain Res 2010;200(1):3–24.

[6] Ajina S, Bridge H. Blindsight and Unconscious Vision: What They Teach Us about the Human Visual System. Neuroscientist 2016.

[7] Das A, Huxlin KR. New approaches to visual rehabilitation for cortical blindness: outcomes and putative mechanisms. Neuroscientist 2010;16(4):374–87.

[8] Shadlen MN, Newsome WT. Motion perception: seeing and deciding. Proc Natl Acad Sci U S A 1996;93(2):628–33.

[9] Britten KH, Shadlen MN, Newsome WT, Movshon JA. Responses of neurons in macaque MT to stochastic motion signals. Vis Neurosci 1993;10(6):1157–69.

[10] Britten KH, Shadlen MN, Newsome WT, Movshon JA. The analysis of visual motion: a comparison of neuronal and psychophysical performance. J Neurosci 1992;12(12):4745–65.

[11] Newsome WT, Pare EB. A selective impairment of motion perception following lesions of the middle temporal visual area (MT). J Neurosci 1988;8(6):2201–11.

[12] Newsome WT, Wurtz RH, Dursteler MR, Mikami A. Deficits in visual motion processing following ibotenic acid lesions of the middle temporal visual area of the macaque monkey. J Neurosci 1985;5(3):825–40.

[13] Zihl J, von Cramon D, Mai N. Selective disturbance of movement vision after bilateral brain damage. Brain 1983;106 (Pt 2):313–40.

[14] Tootell RB, Reppas JB, Dale AM, Look RB, Sereno MI, Malach R, et al. Visual motion aftereffect in human cortical area MT revealed by functional magnetic resonance imaging. Nature 1995;375(6527):139–41.

[15] Salzman CD, Murasugi CM, Britten KH, Newsome WT. Microstimulation in visual area MT: effects on direction discrimination performance. J Neurosci 1992;12(6):2331–55.

[16] Law CT, Gold JI. Neural correlates of perceptual learning in a sensory-motor, but not a sensory, cortical area. Nat Neurosci 2008;11(4):505–13.

[17] Law CT, Gold JI. Shared mechanisms of perceptual learning and decision making. Top Cogn Sci 2010;2(2):226–38.

[18] Law CT, Gold JI. Reinforcement learning can account for associative and perceptual learning on a visual-decision task. Nat Neurosci 2009;12(5):655–63.

[19] Liu LD, Pack CC. The Contribution of Area MT to Visual Motion Perception Depends on Training. Neuron 2017;95(2):436–46 e3.

[20] Larcombe SJ, Kennard C, Bridge H. Increase in MST activity correlates with visual motion learning: A functional MRI study of perceptual learning. Hum Brain Mapp 2017.

[21] Rokem A, Silver MA. Cholinergic enhancement augments magnitude and specificity of visual perceptual learning in healthy humans. Curr Biol 2010;20(19):1723–8.

[22] Antal A, Kincses TZ, Nitsche MA, Paulus W. Modulation of moving phosphene thresholds by transcranial direct current stimulation of V1 in human. Neuropsychologia 2003;41(13):1802–7.

[23] Antal A, Kincses TZ, Nitsche MA, Paulus W. Manipulation of phosphene thresholds by transcranial direct current stimulation in man. Exp Brain Res 2003;150(3):375–8.

[24] Kraft A, Roehmel J, Olma MC, Schmidt S, Irlbacher K, Brandt SA. Transcranial direct current stimulation affects visual perception measured by threshold perimetry. Exp Brain Res 2010;207(3-4):283–90.

[25] Olma MC, Kraft A, Roehmel J, Irlbacher K, Brandt SA. Excitability changes in the visual cortex quantified with signal detection analysis. Restor Neurol Neurosci 2011;29(6):453–61.

[26] Antal A, Nitsche MA, Kruse W, Kincses TZ, Hoffmann KP, Paulus W. Direct current stimulation over V5 enhances visuomotor coordination by improving motion perception in humans. J Cogn Neurosci 2004;16(4):521–7.

[27] Sczesny-Kaiser M, Beckhaus K, Dinse HR, Schwenkreis P, Tegenthoff M, Hoffken O. Repetitive Transcranial Direct Current Stimulation Induced Excitability Changes of Primary Visual Cortex and Visual Learning Effects-A Pilot Study. Front Behav Neurosci 2016;10:116.

[28] Reis J, Schambra HM, Cohen LG, Buch ER, Fritsch B, Zarahn E, et al. Noninvasive cortical stimulation enhances motor skill acquisition over multiple days through an effect on consolidation. Proc Natl Acad Sci U S A 2009;106(5):1590–5.

[29] O’Shea J, Revol P, Cousijn H, Near J, Petitet P, Jacquin-Courtois S, et al. Induced sensorimotor cortex plasticity remediates chronic treatment-resistant visual neglect. Elife 2017;6.

[30] Garcia-Perez MA. Forced-choice staircases with fixed step sizes: asymptotic and small-sample properties. Vision Res 1998;38(12):1861–81.

[31] Hotson JR, Anand S. The selectivity and timing of motion processing in human temporo-parieto-occipital and occipital cortex: a transcranial magnetic stimulation study. Neuropsychologia 1999;37(2):169–79.

[32] Walsh V, Ashbridge E, Cowey A. Cortical plasticity in perceptual learning demonstrated by transcranial magnetic stimulation. Neuropsychologia 1998;36(4):363–7.

[33] Larcombe SJ, Kennard C, Bridge H. Time course influences transfer of visual perceptual learning across spatial location. Vision Res 2017;135:26–33.

[34] Wagenmakers EJ, Love J, Marsman M, Jamil T, Ly A, Verhagen J, et al. Bayesian inference for psychology. Part II: Example applications with JASP. Psychon Bull Rev 2018;25(1):58–76.

[35] Wagenmakers EJ, Marsman M, Jamil T, Ly A, Verhagen J, Love J, et al. Bayesian inference for psychology. Part I: Theoretical advantages and practical ramifications. Psychon Bull Rev 2018;25(1):35–57.

[36] Antal A, Varga ET, Nitsche MA, Chadaide Z, Paulus W, Kovacs G, et al. Direct current stimulation over MT+/V5 modulates motion aftereffect in humans. Neuroreport 2004;15(16):2491–4.

[37] Watson JD, Myers R, Frackowiak RS, Hajnal JV, Woods RP, Mazziotta JC, et al. Area V5 of the human brain: evidence from a combined study using positron emission tomography and magnetic resonance imaging. Cereb Cortex 1993;3(2):79–94.

[38] Malikovic A, Amunts K, Schleicher A, Mohlberg H, Eickhoff SB, Wilms M, et al. Cytoarchitectonic analysis of the human extrastriate cortex in the region of V5/MT+: a probabilistic, stereotaxic map of area hOc5. Cereb Cortex 2007;17(3):562–74.

[39] Dumoulin SO, Bittar RG, Kabani NJ, Baker CL, Jr., Le Goualher G, Bruce Pike G, et al. A new anatomical landmark for reliable identification of human area V5/MT: a quantitative analysis of sulcal patterning. Cereb Cortex 2000;10(5):454–63.

[40] Large I, Bridge H, Ahmed B, Clare S, Kolasinski J, Lam WW, et al. Individual Differences in the Alignment of Structural and Functional Markers of the V5/MT Complex in Primates. Cereb Cortex 2016;26(10):3928–44.

[41] Datta A, Truong D, Minhas P, Parra LC, Bikson M. Inter-Individual Variation during Transcranial Direct Current Stimulation and Normalization of Dose Using MRI-Derived Computational Models. Front Psychiatry 2012;3:91.

[42] Battaglini L, Noventa S, Casco C. Anodal and cathodal electrical stimulation over V5 improves motion perception by signal enhancement and noise reduction. Brain Stimul 2017;10(4):773–9.

[43] Minarik T, Berger B, Althaus L, Bader V, Biebl B, Brotzeller F, et al. The Importance of Sample Size for Reproducibility of tDCS Effects. Front Hum Neurosci 2016;10:453.

[44] Tikhonov A, Haarmeier T, Thier P, Braun C, Lutzenberger W. Neuromagnetic activity in medial parietooccipital cortex reflects the perception of visual motion during eye movements. NeuroImage 2004;21(2):593–600.

